# Coordinated microRNA–mRNA regulatory remodelling across the compressed maturation of the African killifish (*Nothobranchius furzeri*)

**DOI:** 10.64898/2026.09.29.755555

**Authors:** Zainab Afzal, Charles Hatcher, Deepak Kumar

## Abstract

The turquoise killifish (*Nothobranchius furzeri*) is an emerging vertebrate model for development and ageing due to its exceptionally rapid maturation and short lifespan. However, how microRNA (miRNA) regulation coordinates gene expression across its compressed developmental trajectory remains poorly understood. Here, we integrated miRNA and mRNA expression from the same unexposed animals across three developmental stages: newly hatched fry (4 weeks total age), juveniles (∼8 weeks total age), and aged adults (∼8 months of age). Developmental stage was the dominant source of variation in both datasets, with miRNA and mRNA profiles independently resolving a consistent progression from newly hatched fry to juveniles and adults. Transcriptomic changes reflected ongoing tissue maturation, including increased expression of crystallin genes associated with lens development, while more than 2,000 genes were differentially expressed during the juvenile transition. Integration of 398 developmentally dynamic miRNAs with predicted mRNA targets identified specific miRNAs whose developmental changes were significantly enriched for opposing changes in their target genes. These included conserved developmental regulators from the miR-200/141, miR-29, let-7, and miR-34 families, which increased toward adulthood as their predicted target cohorts decreased. Stage-controlled analyses further identified miR-34a, miR-29a, miR-29b, miR-15a, and a novel candidate whose target relationships persisted independently of the overall developmental trajectory. Notably, miR-29 showed particularly strong induction during maturation, consistent with its conserved roles in extracellular-matrix remodeling, differentiation, and vertebrate ageing. Together, these findings reveal coordinated remodeling of conserved miRNA–mRNA regulatory programs across the rapid maturation of *N. furzeri* and establish a framework for investigating post-transcriptional regulation during vertebrate development in a species with limited miRNA annotation.

## Introduction

The African killifish, *Nothobranchius furzeri*, has emerged as a powerful vertebrate model for studying development, regeneration, and ageing within a highly compressed life history. Among vertebrates maintained in captivity, *N. furzeri* has an exceptionally short lifespan and reaches sexual maturity within only a few weeks after hatching^[1–7]^. This rapid progression provides an unusual opportunity to examine major developmental and maturational transitions over a timescale of weeks rather than the months to years required in longer-lived vertebrate models.

During this period, newly hatched fish undergo extensive growth and tissue maturation as proliferative programs transition toward differentiated states and organ systems acquire mature structural and physiological properties. Thus, the compressed maturation of *N. furzeri* provides a tractable system for identifying regulatory mechanisms that coordinate post-embryonic vertebrate development.

MicroRNAs (miRNAs) are small, ∼22-nucleotide non-coding RNAs that provide an important layer of post-transcriptional gene regulation during these developmental transitions^[8–11]^. Through sequence-dependent recognition of target messenger RNAs (mRNAs), miRNAs can reduce transcript stability and/or translation, thereby refining the timing and magnitude of developmental gene expression^[12–14]^. Conserved miRNA families regulate processes including proliferation, differentiation, neurogenesis, myogenesis, epithelial identity, extracellular-matrix remodeling, and tissue maturation. For example, let-7 family members are closely associated with transitions from proliferative to differentiated cellular states^[15, 16]^, the miR-200 family contributes to epithelial differentiation and identity^[17, 18]^, and miR-29 regulates extracellular-matrix and tissue-remodeling programs^[19–21]^. miR-34 family members similarly participate in differentiation, cell-cycle regulation, and tissue homeostasis^[22]^. These regulatory functions make miRNAs particularly relevant to a species in which multiple developmental transitions occur over a remarkably short period.

Importantly, developmental changes in miRNA abundance alone do not establish their regulatory consequences. Because miRNAs primarily act through their target transcripts, understanding their contribution to developmental progression requires integration of miRNA and mRNA expression. A developmental increase in a miRNA accompanied by decreased expression of its predicted target cohort provides stronger evidence of regulatory activity than changes in either molecular layer considered independently. Conversely, miRNA and target genes may change together simply because both respond to developmental stage. Distinguishing these possibilities requires analyses that evaluate anti-correlated miRNA–mRNA relationships while accounting for the shared transcriptional trajectory associated with maturation. Measuring the miRNAome and transcriptome in the same biological individuals therefore provides a particularly powerful framework for identifying candidate regulatory relationships underlying developmental transitions.

Despite the growing use of *N. furzeri* as a vertebrate model, its post-transcriptional regulatory landscape remains comparatively under-characterized. This limitation is particularly evident for miRNAs. *N. furzeri* is not currently represented in miRBase, requiring miRNA identification to rely on sequence conservation with annotated species, such as zebrafish (*Danio rerio*), together with de novo prediction. A species-specific killifish miRNA catalogue was previously generated by Baumgart et al. (2017)^[23]^, providing an important foundation for miRNA annotation in *N. furzeri*. However, coordinated changes between miRNAs and their corresponding mRNA targets across post-hatch maturation have not been systematically characterized. Although the killifish transcriptome is substantially better annotated, it likewise has not been integrated with small-RNA profiles from the same animals to define regulatory networks operating across this rapid developmental interval.

Here, we address this gap by jointly profiling the miRNAome and transcriptome of the same unexposed *N. furzeri* across three stages spanning post-hatch maturation: newly hatched fry at approximately 4 weeks total age, juveniles approximately 4 weeks post-hatch (8 weeks total age), and aged adults (about 8 months old). These stages capture progression from the newly hatched state through the juvenile period in which rapid growth and sexual maturation occur and into adulthood. By sampling this compressed trajectory, we sought to identify coordinated regulatory changes accompanying processes such as proliferation-to-differentiation transitions, extracellular-matrix remodeling, and epithelial, muscular, and other tissue maturation.

To define this regulatory landscape, we first independently characterized developmental variation in small-RNA and mRNA expression and asked whether the two molecular layers resolve a common maturation trajectory. We then identified developmentally dynamic miRNAs and predicted their mRNA targets using *N. furzeri* 3′ untranslated regions. Rather than assuming that sequence-based target predictions represent functional interactions, we tested whether the predicted targets of individual miRNAs were enriched for expression changes opposite to those of their corresponding miRNA. This framework allows regulatory signal to be evaluated at the level of individual miRNAs against an empirical shuffled-target background. We further applied a stage-controlled analysis to distinguish miRNA–target relationships that persist within developmental stages from correlations produced primarily by the shared progression from newly hatched fish to adults. Finally, because *N. furzeri* lacks comprehensive miRNA annotation, we compared newly identified candidates with the previously published species-specific killifish miRNA catalogue to provide independent support for candidate miRNAs.

Using this integrated approach, we identify coordinated remodeling of conserved miRNA– mRNA regulatory modules across killifish maturation. Developmental progression was independently captured by both molecular layers, while integration highlighted conserved miRNA families including miR-200/141, miR-29, let-7, and miR-34 whose increasing expression toward adulthood was accompanied by decreasing expression of predicted target cohorts. Stage-controlled analyses further identified a subset of miRNAs, including miR-34a, miR-29a, miR-29b, and miR-15a, whose target relationships remained detectable after accounting for the overall developmental trajectory. Particularly strong induction of the miR-29 family points to extracellular-matrix and tissue-remodeling regulation as a prominent component of post-hatch maturation. Together, these analyses establish an integrated miRNA–mRNA framework for *N. furzeri* and reveal conserved post-transcriptional regulatory programs associated with its unusually rapid transition from newly hatched fish through juvenile maturation to adulthood.

## Material and Methods

### Animal husbandry

African turquoise killifish (*Nothobranchius furzeri*; ZMZ and GRZ strains) were maintained in a recirculating aquatic system (Aquaneering, San Diego, CA, USA) at 26–27 °C and a water pH of 7.0–7.5. Adult fish were housed in 2.78-L tanks under a 14 h light:10 h dark photoperiod. System water was prepared from reverse-osmosis (RO) water supplemented with Instant Ocean® sea salt. A concentrated salt stock was prepared by dissolving 40 g of sea salt in 1 L of RO water, and 1.5 mL of this stock was added per liter of RO water to achieve a final salt concentration of approximately 60 µg/mL. Adult fish were fed live brine shrimp (Artemia; INVE Aquaculture Inc.) three times daily and supplemented with Otohime C1 fish diet (Reed Mariculture) and frozen bloodworms (Hikari Bio-Pure Blood Worm Cubes; That Fish Place, Cat. No. 284799). Males and females were housed separately, and sex was determined based on secondary sexual characteristics, including differences in coloration and body size. Embryos were collected and maintained according to established *N. furzeri* husbandry protocols with minor modifications (Nath et al., 2023). Embryo medium was prepared using 1 L of RO water supplemented with two Ringer’s tablets (MilliporeSigma, Cat. No. 96724). The medium was sterilized by autoclaving at 121 °C and 15 psi for 20 min, supplemented with 0.002% methylene blue, and maintained at pH 7.0–7.5. Embryos were maintained in covered Petri dishes at 26 °C until hatching. All killifish husbandry and experimental procedures were approved by the Institutional Animal Care and Use Committee (IACUC) at North Carolina Central University (Protocol No. DK10112024) and were conducted in accordance with established animal-care protocols.

### RNA extraction, mRNA sequencing, and small RNA sequencing

For integrated mRNA and miRNA analyses across development, whole-body RNA was collected from Nothobranchius furzeri at three developmental stages: newly hatched fry at 1 day post-hatching (approximately 4 weeks total age; n = 3), juveniles at approximately 4 weeks post-hatching (approximately 8 weeks total age; n = 3), and reproductively mature adults (n = 4; 2 males and 2 females). Each biological replicate consisted of an individual fish that was flash-frozen and processed independently, resulting in a total of 10 biological samples. The same RNA preparation from each individual animal was used for both mRNA and small RNA sequencing, allowing direct integration of mRNA and miRNA expression profiles from matched biological samples.

Whole animals were homogenized in TRIzol Reagent (Invitrogen, Cat. No. 15596018). Newly hatched and juvenile samples were homogenized in 50 µL TRIzol, whereas adult samples were homogenized in 1 mL TRIzol. Total RNA was isolated using the Direct-zol RNA Miniprep Kit (Zymo Research, Cat. No. R2073) according to the manufacturer’s instructions, including on-column DNase treatment. RNA concentration and purity were assessed using a NanoDrop One spectrophotometer (Thermo Fisher Scientific), and all samples exhibited A260/A280 ratios >2.0.

Purified total RNA was submitted to Novogene (Sacramento, CA, USA) for library preparation, quality control, and sequencing. For mRNA sequencing, stranded libraries were generated using the Illumina TruSeq Stranded mRNA Library Preparation Kit and sequenced on an Illumina NovaSeq platform to generate 150-bp paired-end reads, with >40 million reads obtained per sample. Small RNA libraries were prepared from the corresponding RNA samples and sequenced on an Illumina NovaSeq platform using 50-bp single-end reads to capture small non-coding RNA species, including miRNAs. All sequencing libraries underwent quality control prior to downstream analyses and met the required sequencing quality thresholds.

### Small RNA processing and quantification

Small RNA sequencing reads were quality- and adapter-trimmed using fastp with the following parameters: ‘--length_required 18’, ‘--length_limit 35’, ‘--qualified_quality_phred 20’, ‘--unqualified_percent_limit 40’, and ‘--trim_poly_x’. The adapter sequence was removed during processing. Read quality and small RNA composition were subsequently assessed using miRTrace with zebrafish (*Danio rerio*; dre) specified as the reference species.

To maintain consistent feature identity across all developmental stages, reads from all libraries were quantified against a fixed miRNA reference catalogue comprising zebrafish-referenced known miRNAs from miRBase and candidate *N. furzeri* miRNAs identified *de novo* using miRDeep2 against the *N. furzeri* RefSeq genome assembly (GCF_001465895.1; Nfu_20140520). Read counts were assigned using the miRDeep2 ‘quantifier.pl’ workflow. Low-abundance features with fewer than 10 total reads across all samples were excluded prior to downstream analysis.

Count data were analyzed using DESeq2. Variance-stabilizing transformation (VST) was used for visualization and principal component analysis (PCA), while differential expression was performed on raw counts using DESeq2 Wald tests. Developmental-stage comparisons were performed among newly hatched fish (4 weeks total age), juveniles (8 weeks total age), and reproductively mature adults (8 months of age). Differentially expressed miRNAs were identified using a Benjamini–Hochberg false discovery rate (FDR) threshold of <0.05.

### mRNA alignment and quantification

Paired-end mRNA sequencing reads were quality- and adapter-trimmed using fastp with adapter auto-detection and the parameters ‘--detect_adapter_for_pè, ‘--qualified_quality_phred 20’, ‘-- unqualified_percent_limit 40’, and ‘--length_required 36’. Trimmed reads were aligned to the *N. furzeri* RefSeq genome assembly (GCF_001465895.1) using HISAT2. A genome index was generated from the reference assembly, and known splice sites derived from the corresponding RefSeq annotation were supplied during alignment. The ‘--dtà option was used to retain alignments suitable for downstream transcript-level analyses.

Gene-level read counts were generated using featureCounts with exons specified as the counting feature and ‘gene_id’ as the grouping attribute. Paired-end fragments were counted using the ‘-- countReadPairs’ option. Library strandedness was evaluated empirically by comparing featureCounts assignments under alternative strandedness settings, and the libraries were subsequently analyzed as unstranded (‘-s 0’).

Genes with fewer than 10 total reads across all samples were excluded before downstream analysis. Differential gene expression was analyzed using DESeq2 with developmental stage included as the primary model factor and ordered as 4-week newly hatched fry, 8-week juveniles, and 8 month old adults. Raw counts were used for differential expression testing, whereas VST-transformed expression values were used for PCA and other expression-based analyses. PCA was performed using the 2,000 genes with the greatest variance across samples. Pairwise developmental-stage contrasts were evaluated using Wald tests with Benjamini–Hochberg correction, and genes with FDR <0.05 were considered differentially expressed.

### miRNA–mRNA target prediction and integration

To identify potential regulatory relationships between developmentally dynamic miRNAs and mRNAs, miRNAs exhibiting significant differential expression in at least one developmental-stage comparison (FDR <0.05) were subjected to target prediction. Candidate target sites were identified using miRanda against *N. furzeri* 3′ untranslated region (3′UTR) sequences. For each gene, the longest annotated 3′UTR of at least 20 nucleotides was extracted in a strand-aware manner from exon and stop-codon coordinates in the RefSeq annotation. miRanda target prediction was performed using a minimum alignment score of 150, maximum free energy of −20 kcal/mol, and strict seed-matching criteria (‘-sc 150 -en -20 -strict’).

For each predicted miRNA–mRNA pair, developmental expression trajectories were compared across the three stages. miRNA abundance was represented as log2-transformed counts per million (CPM), and mRNA abundance was represented by VST-transformed expression values. Pearson correlation coefficients were calculated across the 4-week, 8-week, and 8 month old adult stage means. Negative correlations were considered consistent with canonical miRNA-mediated repression.

To determine whether the predicted targets of an individual miRNA exhibited anti-correlated expression more frequently than expected by chance, the observed proportion of negatively correlated targets was compared with a miRNA-specific shuffled-target null distribution. For each miRNA, random gene sets equal in size to its predicted target set were sampled with replacement from the filtered mRNA expression dataset and evaluated for anti-correlation with the corresponding miRNA trajectory. This procedure was repeated 100,000 times per miRNA.

Empirical statistics were calculated from the resulting null distributions, and P values were corrected for multiple testing using the Benjamini–Hochberg procedure.

### Stage-controlled miRNA–mRNA correlation analysis

As correlations calculated from three developmental-stage means may reflect a shared developmental trajectory rather than direct regulatory coupling, an additional within-stage analysis was performed. For each miRNA–target pair, miRNA and mRNA expression values were residualized relative to their respective developmental-stage means across the 10 individual animals. Pearson correlations were then calculated between the residualized miRNA and mRNA expression values.

For each candidate miRNA, the mean residual correlation across its predicted targets was compared with a target-shuffled null distribution generated using 2,000 permutations (random seed = 42). A one-sided test was used to identify miRNAs whose targets exhibited stronger negative residual correlations than expected by chance, followed by Benjamini–Hochberg correction. This analysis was used to identify miRNA–target relationships that persisted after accounting for the overall developmental-stage trajectory.

### Sensitivity analyses

Sensitivity analyses were performed to evaluate the robustness of miRNA–mRNA relationships to adult sex and correlation methodology. To assess potential sex-associated effects at the adult stage, target-enrichment analyses were repeated using adult males and adult females separately. To evaluate dependence on the correlation metric, anti-correlated target enrichment was additionally assessed using Spearman rank correlations and correlations calculated across individual animals rather than the three developmental-stage means.

As whole-body RNA profiles may also reflect developmental changes in tissue composition, developmentally dynamic miRNAs were evaluated against canonical vertebrate tissue-associated miRNA families, including muscle-associated myomiRs and hepatic, neural, erythroid/immune, and epithelial miRNAs. Conserved developmental regulatory modules, including miRNAs associated with muscle differentiation, maternal transcript clearance, proliferation-to-differentiation transitions, epithelial identity, and extracellular-matrix remodeling, were additionally examined in the context of their corresponding mRNA expression programs.

### Cross-species conservation and development–ageing annotation

To characterize evolutionary conservation, mature sequences from the miRNA reference catalogue were compared with mature miRNA sequences represented in miRBase through RNAcentral. For *de novo* candidates, the predicted mature consensus sequence was used, whereas known miRNAs were represented by their corresponding miRBase mature sequences. Where necessary, miRNA names were normalized for copy number and mature arm designation.

For each miRNA, the seed sequence was defined as nucleotides 2–7 of the mature sequence. Seed conservation was assessed across species represented in miRBase, including specific evaluation of conservation in human, mouse, *Drosophila*, and *Caenorhabditis elegans*. Based on taxonomic distribution, miRNAs were categorized according to conservation breadth, ranging from broadly conserved miRNAs to vertebrate-, fish-, zebrafish-shared, or lineage-restricted candidates.

To distinguish developmental regulation from previously reported ageing associations, miRNA families were cross-referenced against a curated set of vertebrate and *N. furzeri* ageing-associated miRNAs derived from the published literature, including previously reported killifish brain-ageing signatures. Where available, the reported direction of age-associated expression change was recorded and compared with the direction of expression across maturation in the present dataset. miRNAs were subsequently annotated as developmentally dynamic, previously ageing-associated, associated with both, or neither. For miRNAs represented in both categories, directional concordance between maturation-associated and previously reported ageing-associated changes was assessed.

This literature-based annotation was used solely to place developmentally regulated miRNAs within the context of previously reported ageing-associated pathways. As the present study does not include a longitudinal or cross-sectional adult ageing series, these analyses were not used to identify ageing-specific miRNAs.

## Results

### miRNA and mRNA expression profiles reveal coordinated remodeling across killifish maturation

To define molecular changes across the compressed developmental trajectory of *Nothobranchius furzeri*, we examined miRNA and mRNA expression at three stages spanning early post-hatch development through adulthood (Fig. 1A). Newly hatched fish were collected at approximately 4 weeks total age, older juveniles at approximately 8 weeks total age, and adults at approximately 8 months of age. This design captures the transition from newly hatched fish through a period of rapid juvenile growth and maturation to the adult state.

**Figure 1.**
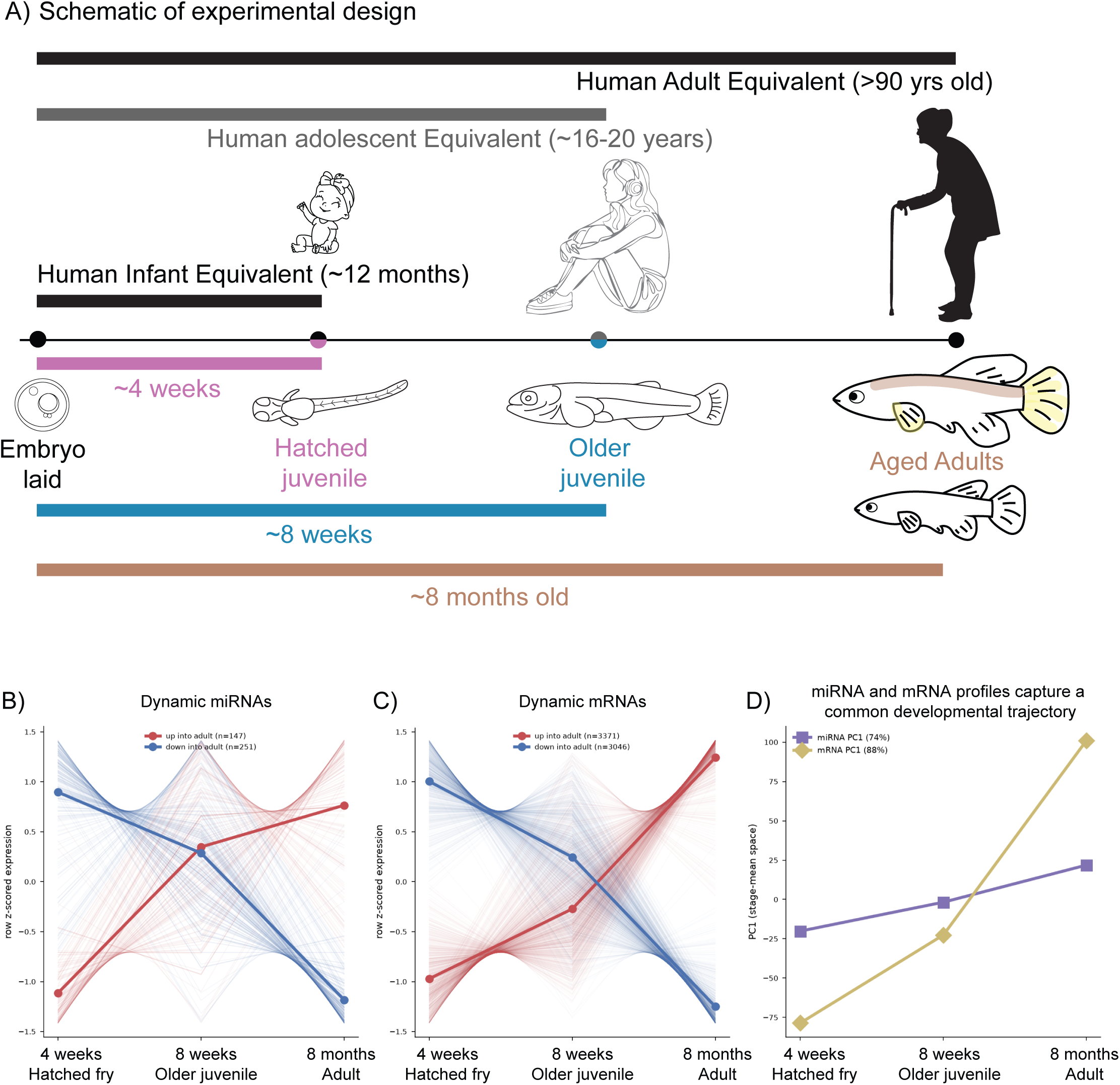
miRNA and mRNA expression profiles reveal coordinated remodeling across killifish maturation. (A) Schematic of the experimental design showing the three *Nothobranchius furzeri* life stages examined: newly hatched fish at approximately 4 weeks total age, older juveniles at approximately 8 weeks total age, and adults at approximately 8 months of age. Whole-body RNA from individual animals was used for matched miRNA and mRNA profiling. (B) Developmental trajectories of dynamically expressed miRNAs across the three stages. Row z-scored expression is shown for individual miRNAs, separated according to whether expression increased (red; n=147) or decreased (blue; n=251) toward adulthood. Thick lines indicate the mean trajectory for each group. (C) Developmental trajectories of dynamically expressed mRNAs, shown as row z-scored expression and separated into transcripts that increased (red; n=3,371) or decreased (blue; n=3,046) toward adulthood. (D) Principal component analysis of stage-level expression profiles demonstrates concordant developmental ordering of the miRNA and mRNA datasets from newly hatched fish to older juveniles and adults. PC1 accounted for 74% of stage-level variation in miRNA expression and 88% of variation in mRNA expression. Together, the two molecular layers independently capture a common expression trajectory across killifish maturation.

Both miRNA and mRNA expression underwent extensive remodeling across this developmental trajectory. Among developmentally dynamic miRNAs, 147 progressively increased and 251 decreased toward adulthood (Fig. 1B). Although individual miRNAs displayed distinct patterns, the overall expression trajectories showed a clear transition across the three stages, with changes emerging between newly hatched and older juvenile fish and becoming more pronounced in adults. Similarly, 3,371 mRNAs increased and 3,046 decreased toward adulthood (Fig. 1C), demonstrating extensive remodeling at both the post-transcriptional and transcriptional levels.

Importantly, the miRNA and mRNA datasets independently captured the same developmental progression. Analysis of the stage-level expression profiles ordered both molecular datasets from newly hatched fish to older juveniles and then adults (Fig. 1D). The first principal component captured 74% of the stage-associated variation in miRNA expression and 88% of the variation in mRNA expression. Thus, despite representing distinct layers of gene regulation, miRNA and mRNA profiles independently identified a common molecular trajectory across maturation.

Together, these findings demonstrate coordinated remodeling of the miRNA and mRNA landscapes across the rapid life history of *N. furzeri*. The agreement between the two molecular layers provided the basis for subsequent analyses examining whether developmental changes in miRNA expression were associated with opposing changes in their predicted mRNA targets.

### Transcriptomic remodeling increases across maturation and supports a progressive developmental transition

The extent of differential expression increased with developmental distance in both the miRNA and mRNA datasets. The batch-matched comparison between newly hatched fish (4 weeks total age) and older juveniles (8 weeks total age) identified 154 differentially expressed miRNAs and 2,129 differentially expressed genes (FDR<0.05). In comparison, 246 miRNAs and 7,502 genes differed between older juveniles and adults, while the largest differences were observed between newly hatched fish and adults, with 389 miRNAs and 9,260 genes differentially expressed (FDR<0.05). Together, these patterns demonstrate that substantial molecular remodeling is already evident during the early post-hatch juvenile transition and becomes progressively more pronounced across maturation.

The mRNA dataset also identified developmental changes consistent with tissue maturation. Among the most strongly induced transcripts in adults were the crystallin genes *crybb3*, *crybb2*, and *crygs*, which encode structural components of the vertebrate lens. *Crybb3* showed particularly strong induction, increasing by 8.06 log2-fold relative to newly hatched fish (FDR=3×10 ¹¹ ). The progressive increase in crystallin expression provides an example of a tissue-associated transcriptional program accompanying the transition toward the adult state.

Importantly, extensive differential expression was already evident between newly hatched fish and older juveniles in both the miRNA and mRNA datasets, demonstrating that substantial molecular remodeling occurs during the juvenile interval and precedes the adult stage. The further increase in differential expression across the juvenile-to-adult and newly hatched-to-adult comparisons supports a progressive reorganization of the regulatory landscape across post-hatch maturation. Together, the coordinated changes observed across both molecular layers indicate that rapid killifish maturation is accompanied by extensive remodeling of miRNA and mRNA expression programs.

### Developmentally regulated miRNAs show repression-consistent changes in predicted mRNA targets

To determine whether changes in miRNA expression across maturation were associated with corresponding changes in their predicted mRNA targets, we integrated the miRNA and mRNA datasets and examined target expression across developmental stages. Across the complete predicted network, approximately half of miRNA–target pairs were anti-correlated (50.5%; mean trajectory *r*=−0.006), indicating that anti-correlation was not a general feature of sequence-predicted targets. Instead, enrichment was observed for specific developmentally regulated miRNAs.

Nine miRNAs showed significant enrichment of anti-correlated predicted targets at FDR<0.05, with an additional set identified at the exploratory threshold of FDR<0.10 (Fig. 2A). Several of the strongest signals belonged to conserved miRNA families associated with developmental transitions. miR-141-3p and miR-200a-3p increased toward adulthood, with 63% and 65% of their predicted targets changing in the opposite direction, respectively. Similarly, miR-29a-3p and miR-29b-3p increased strongly across maturation, with 62% and 60% of their predicted targets showing anti-correlated expression (Table 2). Let-7b-5p also increased toward adulthood, with 60% of its predicted targets exhibiting an opposing developmental trajectory. Comparison with shuffled target sets showed that the observed anti-correlation for these miRNAs exceeded that expected from randomly selected genes (Fig. 2B), with particularly strong enrichment for the miR-200/141 and miR-29 families.

**Figure 2.**
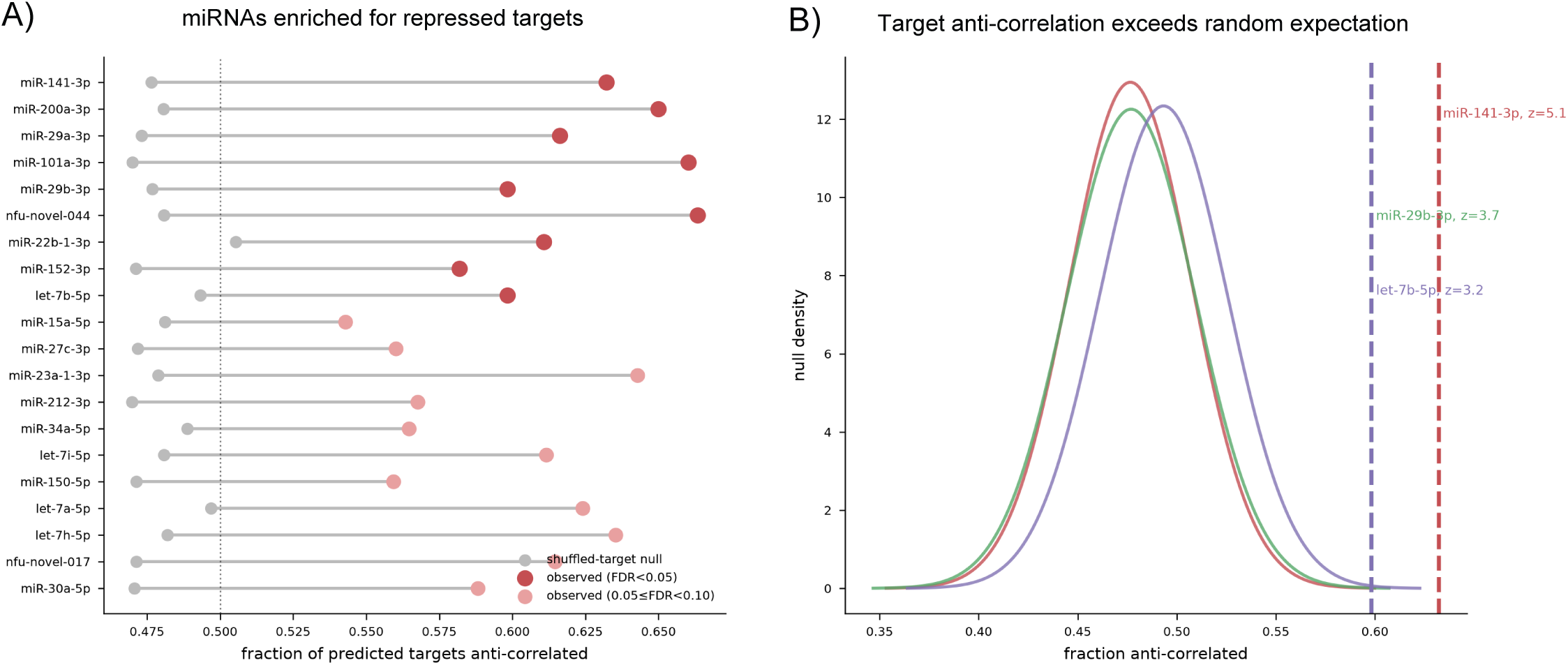
Developmentally regulated miRNAs are enriched for anti-correlated predicted mRNA targets. (A) Fraction of predicted mRNA targets showing expression trajectories anti-correlated with each developmentally regulated miRNA across newly hatched fish, older juveniles, and adults. Colored points indicate the observed fraction of anti-correlated predicted targets for each miRNA, and gray points indicate the expected fraction derived from shuffled target sets. Dark red indicates miRNAs significantly enriched for anti-correlated targets at FDR<0.05, and light red indicates miRNAs meeting the exploratory threshold of 0.05≤FDR<0.10. Dashed vertical line indicates an anti-correlated fraction of 0.5. (B) Representative shuffled-target null distributions for miR-141-3p, miR-29b-3p, and let-7b-5p. Solid curves show the distributions of anti-correlated target fractions obtained from 100,000 randomly generated target sets for each miRNA. Dashed vertical lines indicate the observed anti-correlated target fraction for the corresponding predicted target cohort. The displacement of the observed values from their respective null distributions demonstrates enrichment of repression-consistent target expression beyond that expected from randomly selected genes. FDR values were calculated using Benjamini–Hochberg correction.

**Table 1.** Differential miRNA and mRNA expression increases across killifish maturation. Numbers of differentially expressed miRNAs and mRNAs identified for each pairwise developmental-stage comparison. Differential expression was determined using DESeq2 with Benjamini–Hochberg correction for multiple testing (FDR<0.05). Comparisons represent newly hatched fish (4 weeks total age), older juveniles (8 weeks total age), and 8-month-old adults. FDR, false discovery rate.

| Developmental comparison | Differentially expressed miRNAs | Differentially expressed mRNAs |
| --- | --- | --- |
| 8 weeks vs. 4 weeks | 154 | 2,129 |
| Adult vs. 8 weeks | 246 | 7,502 |
| Adult vs. 4 weeks | 389 | 9,260 |

**Table 2.** Developmentally regulated miRNAs enriched for anti-correlated predicted mRNA targets. Selected developmentally regulated miRNAs showing enrichment of anti-correlated predicted mRNA targets across killifish maturation. Adult versus 4-week log FC represents the change in miRNA expression between newly hatched fish and 8-month-old adults. The anti-correlated target fraction represents the proportion of predicted targets whose developmental expression trajectory was negatively correlated with that of the corresponding miRNA. Enrichment z-scores and FDR values were calculated relative to miRNA-specific shuffled-target distributions. FDR<0.05 was considered significant, while miRNAs with 0.05≤FDR<0.10 were included as an exploratory set for subsequent stage-controlled analyses. ECM, extracellular matrix; FDR, Benjamini–Hochberg false discovery rate.

| miRNA | Developmental module | Adult vs. 4-week $\log_2$FC | Predicted targets (n) | Anti-correlated targets | Enrichment z-score | FDR |
| --- | --- | --- | --- | --- | --- | --- |
| **miR-141-3p** | Epithelial identity | +2.45 | 261 | 0.63 | 5.05 | 0.0020 |
| **miR-200a-3p** | Epithelial identity | +2.22 | 220 | 0.65 | 5.01 | 0.0020 |
| **miR-29a-3p** | ECM remodeling | +5.81 | 198 | 0.62 | 4.04 | 0.0080 |
| **miR-29b-3p** | ECM remodeling | +7.86 | 234 | 0.60 | 3.73 | 0.0099 |
| **let-7b-5p** | Proliferation-to-differentiation transition | +5.61 | 239 | 0.60 | 3.25 | 0.037 |
| **miR-34a-5p** | Proliferation | +2.39 | 379 | 0.56 | 2.95 | 0.054 |
| **let-7a-5p** | Proliferation-to-differentiation transition | +3.76 | 125 | 0.62 | 2.84 | 0.067 |

Individual predicted relationships included let-7b with *smarcc1* and *pcsk9*, miR-29a/b with *tril*, and miR-200a/141 with the shared predicted targets *rab4a* and *cep170* (Supp Table S1). These miRNA–mRNA pairs represent candidate regulatory relationships supported by sequence-based target prediction and opposing developmental expression patterns rather than experimentally validated direct interactions.

Opposing expression across developmental stages, however, can also arise when miRNAs and mRNAs independently respond to the same developmental transition. We therefore tested whether the identified relationships remained detectable after accounting for the shared developmental trajectory. Five miRNAs—miR-34a, miR-29a, miR-29b, miR-15a and the novel candidate nfu-novel-017—retained significant negative associations with their predicted target cohorts after stage adjustment (FDR<0.05; Supplementary Fig. S1A). miR-29a and miR-29b showed particularly strong stage-adjusted target associations. Comparison of across-stage enrichment with within-stage coupling further separated these five miRNAs from relationships that were primarily driven by developmental stage (Supplementary Fig. S1B).

In contrast, miR-200a, miR-141 and let-7b showed strong target anti-correlation across developmental stages but did not retain negative associations after developmental stage was accounted for. These miRNAs therefore appear to participate primarily in stage-coordinated developmental programs, whereas miR-34a, miR-29a, miR-29b and miR-15a showed additional evidence of repression-consistent target relationships within developmental stages.

Analysis of target expression distributions further supported these relationships (Supplementary Fig. 1). Predicted target cohorts associated with the stage-adjusted miRNAs were shifted toward lower expression in adults relative to newly hatched fish compared with the genome-wide background (Supplementary Fig. 2A). Median target expression changes were significantly lower than background for miR-34a, miR-29a, miR-29b and miR-15a (one-sided Mann– Whitney, *P*≤0.002; Supplementary Fig. 2B). nfu-novel-017 showed a similar directional shift but did not reach significance in this analysis.

Together, these analyses identify distinct patterns of miRNA–mRNA coordination during killifish maturation. Some miRNAs, including miR-200a/141 and let-7b, showed target anti-correlation primarily as part of the broader developmental trajectory, whereas a more restricted group—including miR-29a/b, miR-34a and miR-15a—retained repression-consistent target relationships after developmental stage was accounted for. These findings highlight specific miRNA–mRNA modules associated with the regulatory remodeling that accompanies post-hatch maturation.

### Conserved and newly identified miRNA programs show distinct trajectories across maturation

Having identified miRNA families with enriched anti-correlated target cohorts, we next examined how representative regulatory modules changed across the developmental trajectory (Fig. 3A). miR-200/141, miR-29, let-7 and miR-34 family members showed distinct stage-dependent trajectories accompanied by opposing changes in their predicted target cohorts.

**Figure 3.**
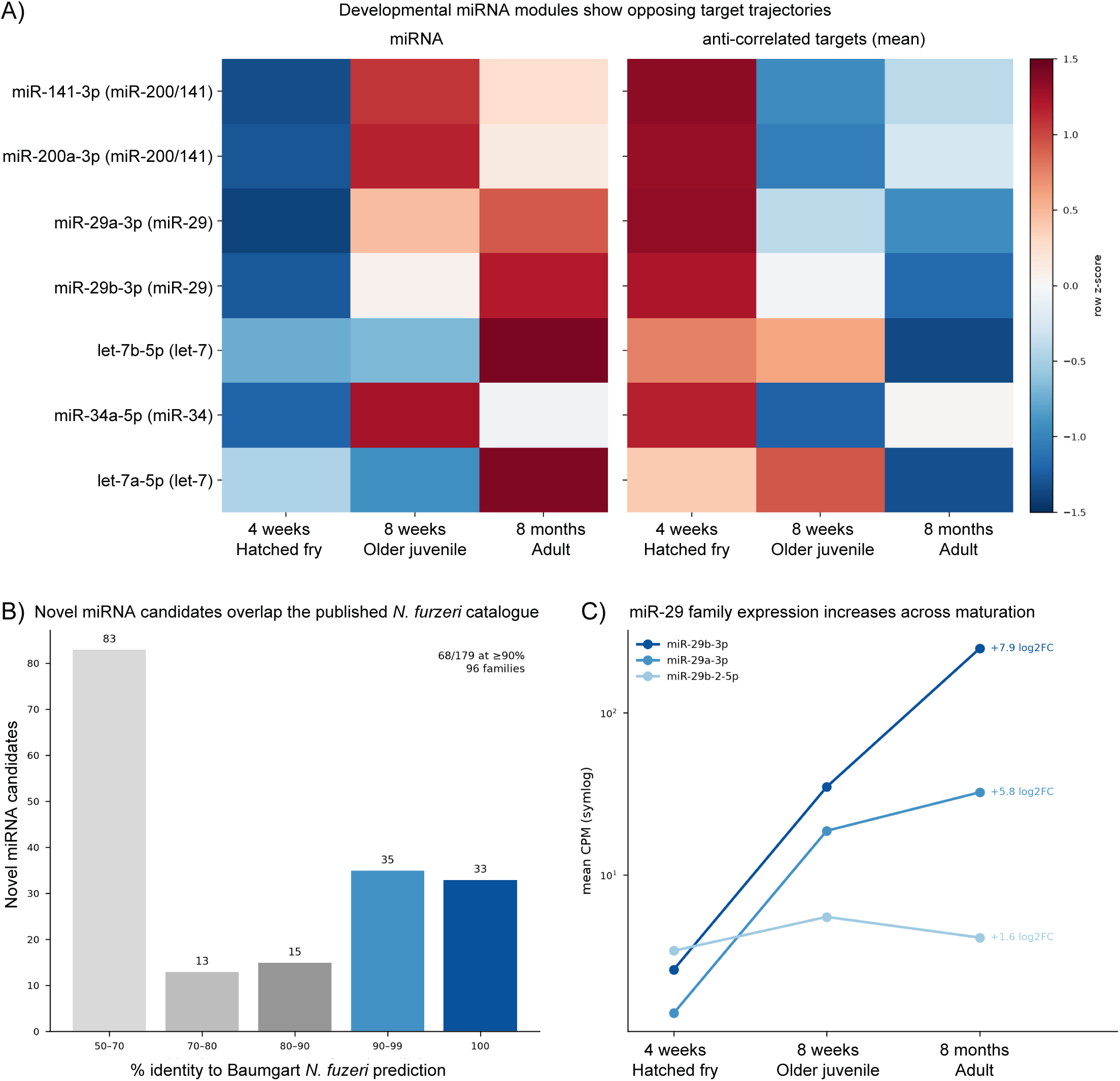
Conserved and newly identified miRNA programs show distinct trajectories across killifish maturation. (A) Expression trajectories of representative developmentally regulated miRNAs (left) and their anti-correlated predicted mRNA target cohorts (right) across newly hatched fish (4 weeks total age), older juveniles (8 weeks total age), and adults (8 months). Values represent row-scaled z-scores of mean expression at each developmental stage. Representative miRNAs from the miR-200/141, miR-29, let-7 and miR-34 families show opposing trajectories between miRNA expression and their predicted target cohorts. (B) Developmental expression trajectories of miR-29 family members. Mean CPM is shown on a logarithmic scale across the three developmental stages. miR-29b-3p and miR-29a-3p increased by +7.9 and +5.8 log FC, respectively, between newly hatched fish and adults, whereas miR-29b-2-5p showed a smaller change (+1.6 log FC). (C) Sequence similarity between de novo N. furzeri miRNA candidates identified in this study and previously predicted N. furzeri miRNAs from Baumgart et al. (2017). Bars show the number of matched candidates within each sequence-identity interval. Of 179 candidates overlapping the published catalogue, 68 showed ≥90% sequence identity, including 33 with 100% identity, representing 96 miRNA families.

Across the seven representative miRNAs shown, adult miRNA expression was increased overall (mean row z-score=+0.75), whereas their anti-correlated target cohorts showed the opposite pattern (mean row z-score=−0.77). These patterns illustrate the coordinated remodeling of representative miRNA–target modules across post-hatch maturation.

The miR-29 family showed one of the most pronounced developmental increases among conserved miRNAs (Fig. 3B). miR-29b-3p increased by 7.86 log -fold from newly hatched fish to adults, from 2.6 to 249.5 mean CPM (FDR=5×10 ¹), while miR-29a-3p increased by 5.81 log -fold, from 1.4 to 32.3 mean CPM (FDR=1×10 ³ ). In contrast, miR-29b-2-5p showed a smaller overall increase (+1.6 log -fold), indicating distinct developmental trajectories among members of the same family. Notably, miR-29a-3p and miR-29b-3p also retained negative associations with their predicted target cohorts after developmental stage was accounted for, highlighting miR-29 as a prominent regulatory module associated with post-hatch maturation.

Other developmentally relevant miRNAs showed complementary trajectories, including increased expression of muscle-associated miR-1 and miR-133 family members toward adulthood and decreased expression of miR-430a-3p from its highest level in newly hatched fish.

To independently assess the de novo miRNA catalogue, we compared the 407 newly identified candidates with the previously published *N. furzeri* miRNA catalogue of Baumgart et al. (2017). A total of 179 candidates matched a previously predicted *N. furzeri* miRNA, including 68 with ≥90% sequence identity and 33 with 100% identity, representing 96 miRNA families (Fig. 3C; Supplementary Table S2). This independent sequence-level overlap supports a substantial subset of the de novo candidates while extending existing *N. furzeri* miRNA resources with expression profiles spanning post-hatch maturation.

### Anti-correlated miRNA target cohorts are associated with distinct developmental pathways

To determine whether anti-correlated target cohorts were associated with distinct biological functions, we examined pathway enrichment for each developmental miRNA module (Fig. 4; Supplementary Table S3). The resulting profiles differed among miRNA families, linking their opposing miRNA–target trajectories to pathways involved in signaling, tissue organization, metabolism and differentiation.

**Figure 4.**
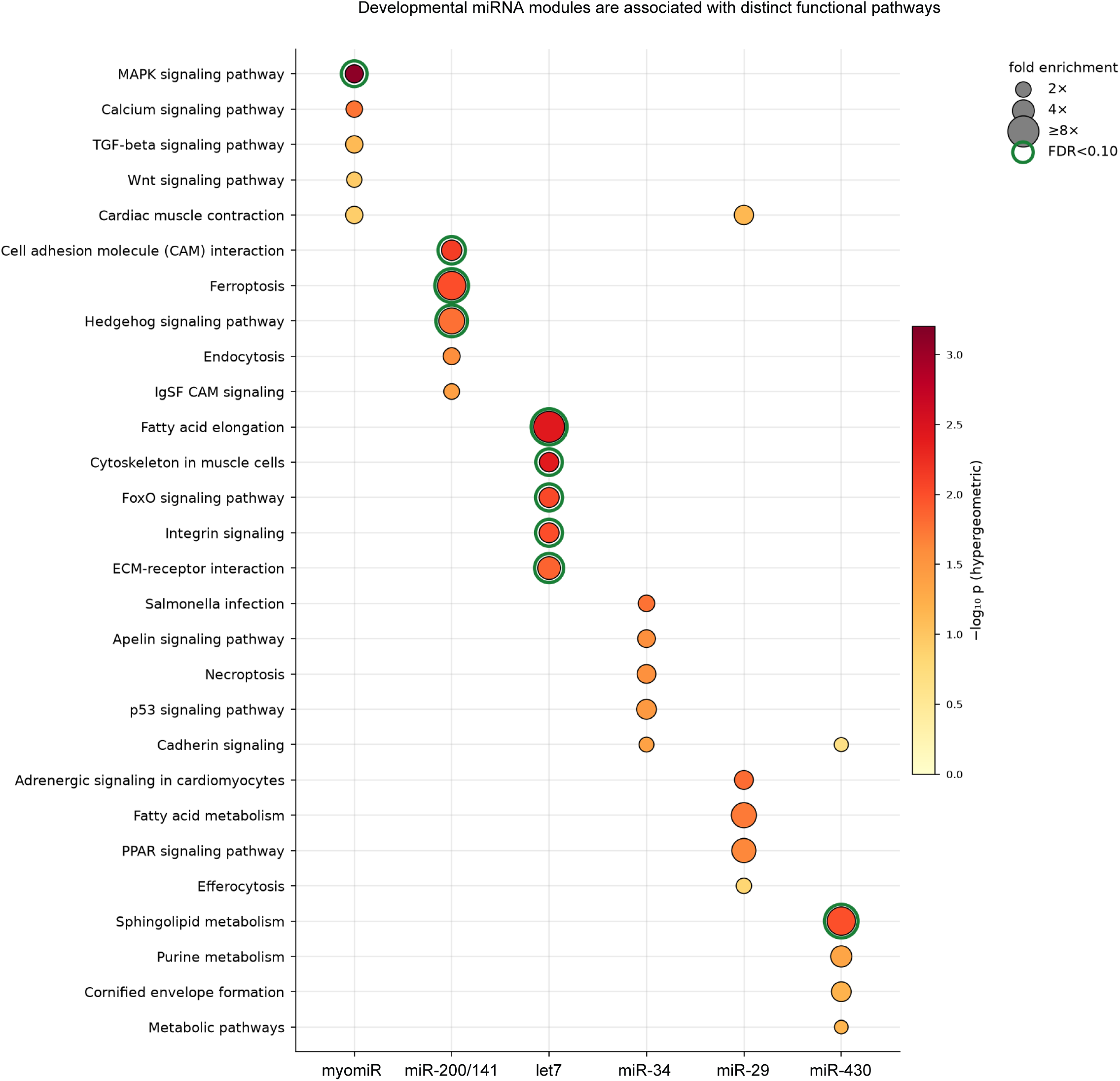
Developmental miRNA modules are associated with distinct functional pathways. KEGG pathway enrichment of anti-correlated predicted mRNA target cohorts for representative developmental miRNA modules. Enrichment was assessed against genes that were both 3′UTR-scannable and KEGG-annotated. Columns represent miRNA modules and rows represent enriched KEGG pathways. Circle size indicates fold enrichment, and color represents enrichment significance as −log (*P*). Green outlines indicate pathways meeting an FDR<0.10 after Benjamini–Hochberg correction. Target cohorts showed module-specific functional associations, including MAPK signaling for myomiRs; cell adhesion molecule interactions, ferroptosis and Hedgehog signaling for miR-200/141; fatty acid elongation, cytoskeletal, FoxO, integrin and ECM-associated pathways for let-7; and sphingolipid metabolism for miR-430. Complete pathway-enrichment results are provided in Supplementary Table S3.

The muscle-associated myomiR target cohort showed significant enrichment for MAPK signaling (13 genes; 2.72-fold enrichment, FDR=0.017), with additional nominal enrichment for calcium, TGF-β and Wnt signaling and cardiac muscle contraction (Fig. 4). The let-7 target cohort showed a broader set of enriched pathways, including fatty acid elongation and cytoskeleton in muscle cells (FDR=0.046), together with FoxO signaling, integrin signaling, ECM–receptor interaction, focal adhesion and calcium signaling (FDR=0.055). These enrichments associate the developmental increase in let-7 expression with coordinated changes in target genes involved in cellular architecture, signaling and tissue remodeling.

The miR-200/141 target cohort was enriched for cell adhesion molecule interactions (3.45-fold, nominal *P*=0.007, FDR=0.089), as well as ferroptosis and Hedgehog signaling at FDR<0.10 (Fig. 4). These enrichments associate the miR-200/141 target cohort with pathways involved in cell interactions and developmental signaling during maturation. The miR-430 cohort similarly showed enrichment for sphingolipid metabolism at FDR<0.10 (6.63-fold, FDR=0.091).

Other modules showed more modest pathway-level enrichment. The miR-34 target cohort included p53 signaling among its nominally enriched pathways (3.25-fold, *P*=0.034), although this association did not remain significant after multiple-testing correction (FDR=0.326). miR-29 targets showed nominal enrichment for adrenergic signaling in cardiomyocytes, fatty acid metabolism and PPAR signaling, but none reached FDR<0.10. Thus, although miR-29 showed strong developmental induction and repression-consistent target associations at the gene level, these targets were not concentrated within a single KEGG pathway under the current annotation.

Together, these results show that anti-correlated targets of developmentally regulated miRNAs are distributed across distinct functional programs, with the strongest pathway-level associations involving signaling, cellular architecture, adhesion and metabolism during post-hatch maturation.

### miRNA–target relationships are robust across analytical approaches and adult sex

We next examined whether the identified miRNA–target relationships were sensitive to adult sex, correlation method, or changes in tissue composition inherent to whole-animal sampling (Fig. 5). Repeating the target-enrichment analysis using either male or female adults retained 19 of the 20 discovery-set miRNAs at FDR<0.10 in each analysis, compared with 20 in the combined analysis (Fig. 5A). The single miRNA not retained differed between the sex-specific analyses, with let-7h absent from the male-only analysis and miR-152 absent from the female-only analysis. Thus, the major target-enrichment patterns were maintained when either adult sex was analyzed separately.

**Figure 5.**
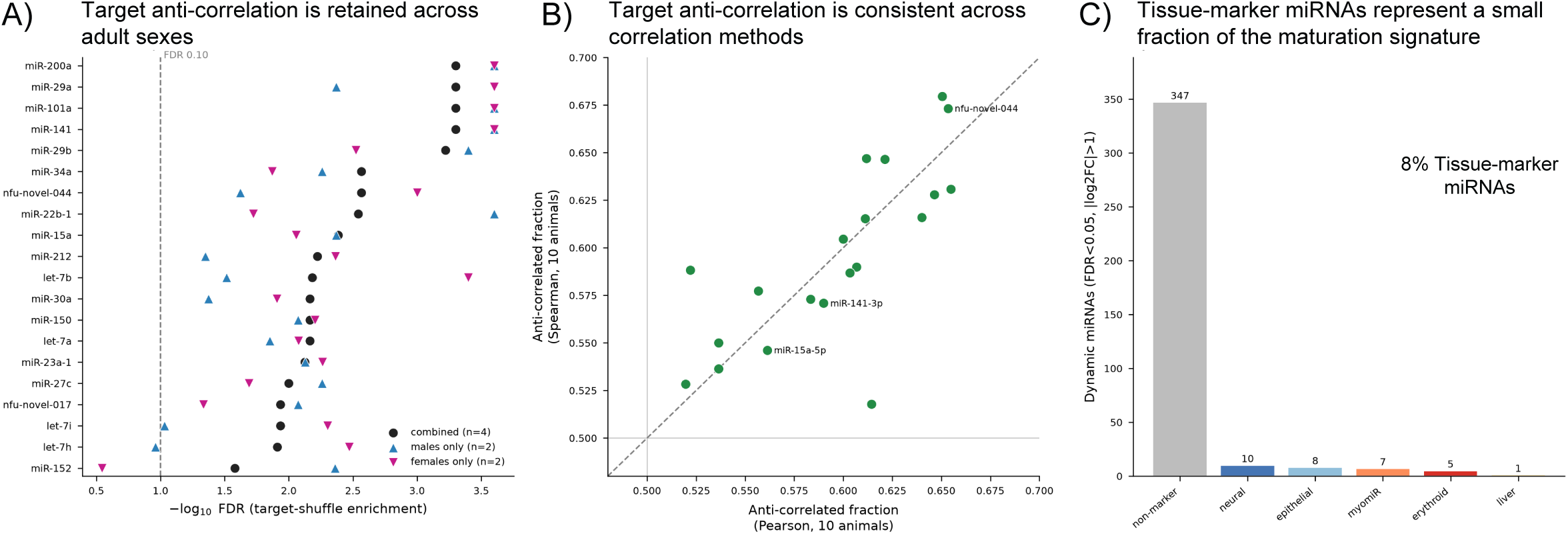
miRNA–target relationships are robust across analytical approaches and adult sex. (A) Sensitivity of miRNA target anti-correlation enrichment to adult sex. Target-enrichment analyses were repeated using all adults (combined; n=4), male adults only (n=2), or female adults only (n=2). Points show −log FDR for enrichment of anti-correlated predicted targets for the 20 miRNAs identified at the discovery threshold (FDR<0.10) in the combined analysis. The dashed vertical line indicates FDR=0.10. Nineteen of the 20 miRNAs remained below the FDR<0.10 threshold in each sex-specific analysis. (B) Comparison of anti-correlated target fractions calculated across individual animals using Pearson and Spearman correlations. Each point represents one miRNA from the discovery set; the dashed diagonal indicates equal values between correlation methods, and horizontal and vertical reference lines indicate an anti-correlated fraction of 0.5. (C) Distribution of canonical tissue-marker annotations among developmentally dynamic miRNAs (FDR<0.05, |log FC|>1). Of the 378 dynamic miRNAs evaluated, 347 were not classified as canonical tissue markers, whereas the remaining miRNAs were assigned to neural (n=10), epithelial (n=8), myomiR/muscle (n=7), erythroid (n=5), or hepatic (n=1) categories. Tissue-associated miRNAs therefore comprised approximately 8% of the dynamic miRNA set.

The anti-correlated target relationships were also consistent across alternative measures of expression association. All 20 enriched miRNAs retained anti-correlated target fractions above 0.5 when calculated using stage-mean Pearson correlations, individual-animal Pearson correlations, or individual-animal Spearman correlations (Fig. 5B). The agreement between individual-level Pearson and Spearman estimates further indicates that the observed relationships were not dependent on a single correlation metric or exclusively on stage-averaged expression patterns.

As the transcriptomic measurements were obtained from whole animals, we additionally examined the contribution of canonical tissue-associated miRNAs to the developmental signature. Of the 378 dynamic miRNAs evaluated in this analysis, 8.2% were classified as canonical tissue-associated markers, with the majority not assigned to a tissue-marker category (Fig. 5C). Tissue-associated miRNAs included muscle, erythroid, epithelial, neural and hepatic markers and showed changes in both directions across maturation. Thus, tissue-associated miRNAs represent a detectable but relatively small component of the broader developmental miRNA signature.

Together, these analyses show that the principal miRNA–target relationships are maintained across adult-sex subsets and alternative correlation approaches, while highlighting tissue composition as an important consideration when interpreting whole-organism developmental expression profiles.

### Developmentally regulated miRNAs include deeply conserved and lineage-restricted regulatory programs

To place the developmental miRNA repertoire in an evolutionary context, we examined seed conservation across 170 taxa represented in miRBase and RNAcentral (Fig. 6A,B). As expected, known zebrafish-referenced miRNAs were dominated by deeply conserved seeds, with 341 of 412 sharing their seed sequence with at least one mammalian miRNA. The *de novo N. furzeri* candidates showed greater diversity in conservation breadth. Of the 407 candidates, 216 contained seeds shared with mammalian miRNAs, 30 showed conservation within other vertebrate or fish groups, and 160 were classified as lineage-restricted. Thus, the de novo catalogue includes both previously unannotated members of conserved miRNA families and candidate killifish-specific regulatory sequences.

**Figure 6.**
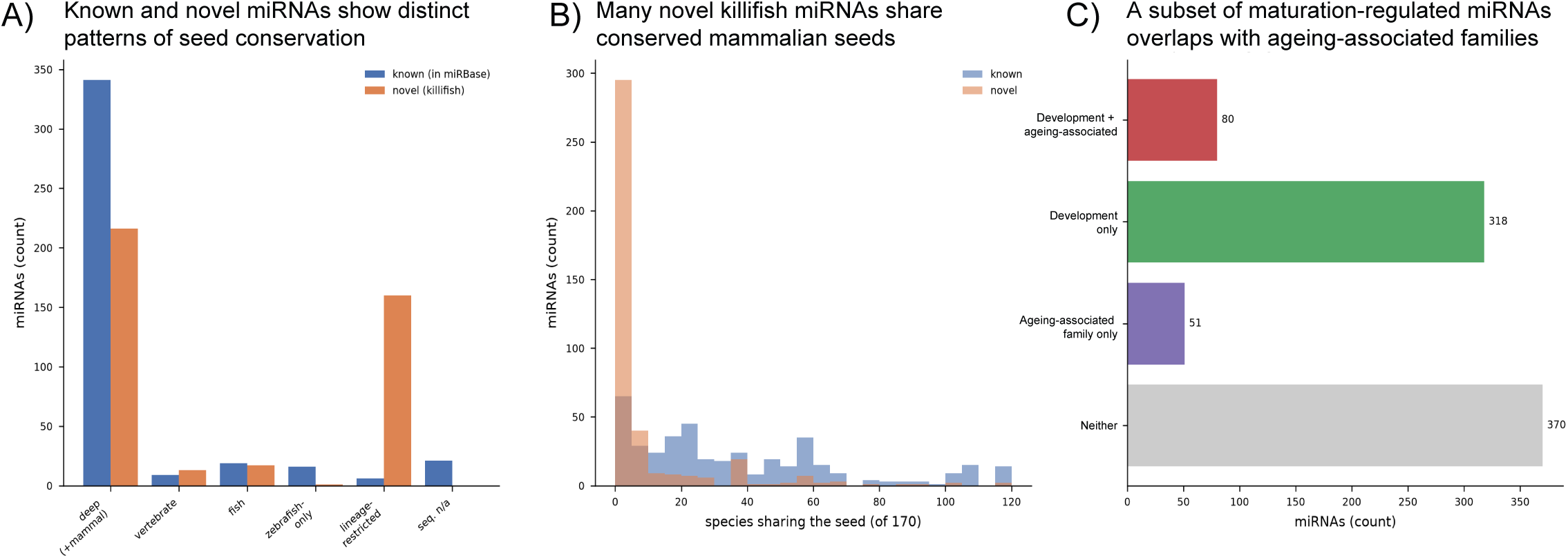
Evolutionary conservation and ageing-associated annotation of the killifish miRNA catalogue. (A) Evolutionary breadth of miRNA seed conservation across 170 taxa represented in the reference databases. The number of species sharing each seed was determined separately for known zebrafish-referenced miRNAs and *N. furzeri* novel candidates. Known miRNAs were predominantly represented by broadly conserved seeds, whereas novel candidates included both conserved and lineage-restricted sequences. (B) Distribution of the number of species sharing each miRNA seed. Of the 407 novel *N. furzeri* candidates, 216 shared a seed with at least one mammalian miRNA, indicating that a substantial proportion of candidates classified as novel in killifish belong to evolutionarily conserved seed families despite the absence of *N. furzeri* from miRBase. (C) Cross-reference of the miRNA catalogue with developmentally dynamic miRNAs identified in this study and miRNA families previously associated with ageing. Among the 819 miRNAs, 318 were developmentally dynamic without an ageing-family annotation, 80 were both developmentally dynamic and members of an ageing-associated family, 51 belonged to ageing-associated families but were not dynamic across the sampled maturation stages, and 370 belonged to neither category. Ageing classifications represent a literature-based annotation and do not indicate ageing-associated expression changes measured directly in this study.

Among the de novo candidates with mammalian seed conservation, conservation breadth varied substantially, ranging from seeds shared by relatively few taxa to those represented broadly across the 170-species reference set (Fig. 6B). This pattern indicates that absence of an existing *N. furzeri* miRBase annotation does not necessarily reflect evolutionary novelty, as many of the newly identified candidates share seed sequences with established vertebrate miRNAs.

We next cross-referenced developmental dynamics in the present dataset with miRNA families previously associated with ageing (Fig. 6C). Most developmentally dynamic miRNAs had no ageing annotation (318 miRNAs), whereas 80 were both developmentally dynamic and members of previously reported ageing-associated families. An additional 51 ageing-associated miRNAs were not significantly dynamic across the developmental stages examined. Among the 80 overlapping miRNAs, 55 changed in the same direction during maturation as previously reported in ageing datasets, whereas 25 showed opposing directional relationships. Thus, a subset of miRNA families shows shared directional changes across maturation and previously reported ageing trajectories, while most developmental changes identified here remain distinct.

Importantly, the present study examines a maturation series rather than an adult ageing series; this comparison represents a literature-based cross-reference rather than a direct measurement of ageing-associated regulation.

Collectively, these analyses define coordinated remodeling of transcriptional and post-transcriptional regulation across rapid post-hatch maturation in *N. furzeri* (Fig. 7). Matched miRNA and mRNA profiling identified extensive developmental changes across both regulatory layers and revealed specific miRNA families associated with repression-consistent changes in predicted target cohorts. These relationships converged on developmental processes involving signaling, cell adhesion, tissue architecture and metabolism and included both deeply conserved and lineage-restricted regulatory programs. Together, the integrated dataset defines a regulatory landscape in which distinct miRNA–mRNA modules are progressively reorganized across the compressed maturation of *N. furzeri*.

**Figure 7.**
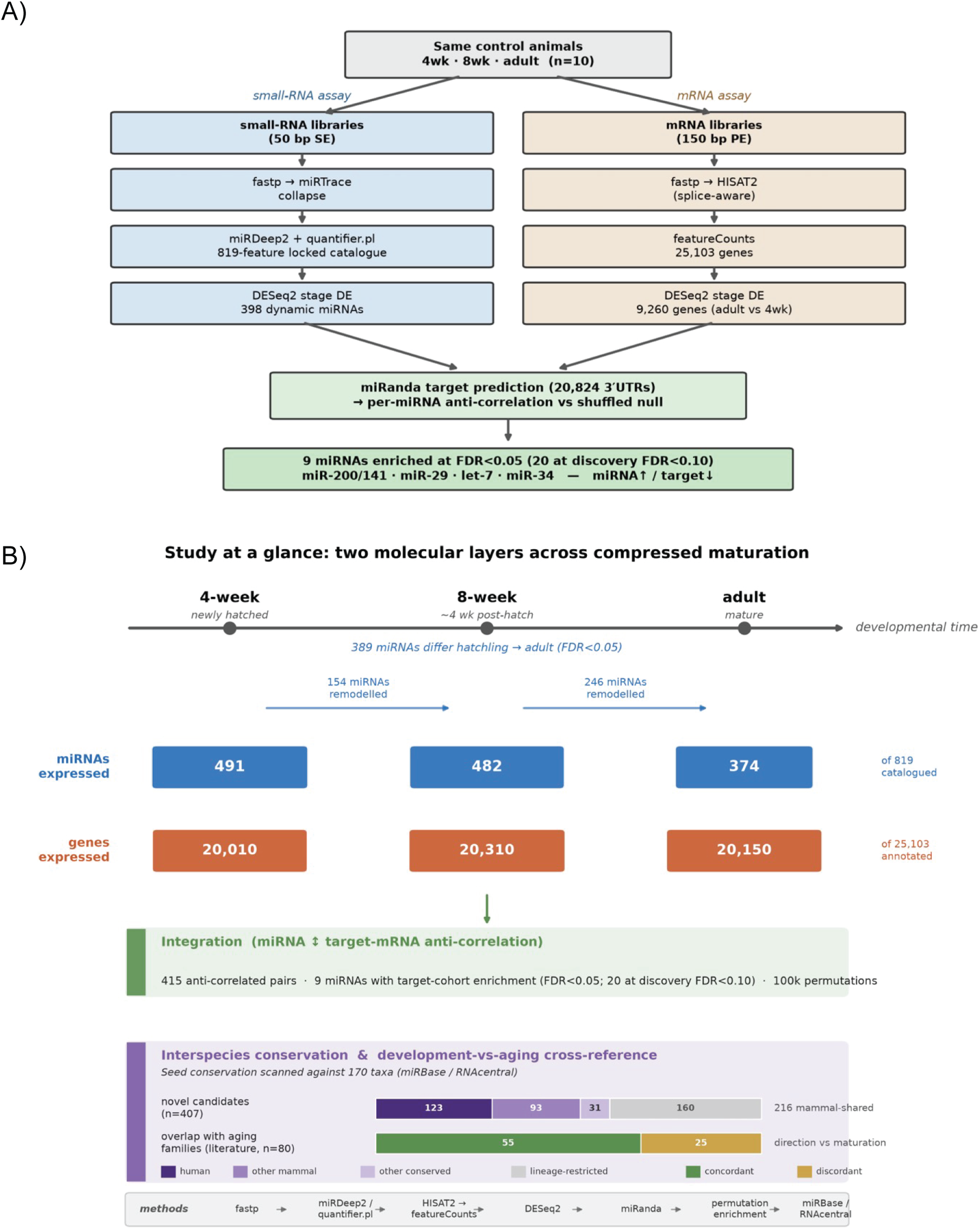
Integrated miRNA and mRNA profiling defines regulatory remodeling across rapid killifish maturation. (A) Overview of the matched small-RNA and mRNA analysis workflow. Small-RNA and mRNA libraries were generated from the same individual N. furzeri sampled at newly hatched (4 weeks total age), older juvenile (8 weeks total age), and adult stages (n=10 total). Small-RNA profiling generated a locked catalogue of 819 miRNA features, of which 398 were developmentally dynamic, while mRNA profiling identified 25,103 annotated genes and 9,260 differentially expressed genes between newly hatched fish and adults. Integration of the two molecular layers using predicted 3′UTR interactions and developmental expression trajectories identified nine miRNAs significantly enriched for anti-correlated predicted targets at FDR<0.05, expanding to 20 miRNAs at the discovery threshold of FDR<0.10. (B) Summary of molecular remodeling across the compressed post-hatch maturation trajectory. Numbers within blue and orange boxes indicate miRNAs and mRNAs expressed at each developmental stage, respectively. Differential expression identified 154 miRNAs remodeled between newly hatched fish and older juveniles, 246 between older juveniles and adults, and 389 across the complete newly hatched-to-adult transition. Integrated miRNA– mRNA analysis identified 415 anti-correlated predicted pairs and specific miRNAs with significant enrichment of anti-correlated target cohorts. Conservation analysis of the 407 de novo miRNA candidates identified both broadly conserved and lineage-restricted seeds, while literature cross-referencing identified 80 developmentally dynamic miRNAs belonging to previously reported ageing-associated families, of which 55 changed in the same direction during maturation as reported in ageing studies. Ageing annotations represent a literature-based cross-reference and not ageing-associated changes measured directly in the present developmental series.

## Discussion

Matched miRNA and mRNA profiling across post-hatch *Nothobranchius furzeri* development revealed coordinated remodeling of transcriptional and post-transcriptional regulation during rapid maturation. Developmental stage was the dominant source of variation in both molecular layers, with progressively greater differences from newly hatched fish to older juveniles and adults. Integration of the datasets further identified specific miRNA families—including miR-200/141, miR-29, let-7 and miR-34—whose developmental trajectories were accompanied by opposing changes in predicted mRNA targets. Rather than anti-correlation being a general property of predicted targets, enrichment was concentrated within specific miRNA families, supporting coordinated regulation of distinct developmental programs.

The identity of these miRNA families is consistent with established roles in developmental transitions^[11]^. Let-7 is a deeply conserved regulator of developmental timing and differentiation^[15, 16]^, while the miR-200 family contributes to epithelial identity and cell-state transitions^[17, 18]^. miR-34 has similarly been linked to p53-dependent control of proliferation and differentiation^[24, 25]^. Particularly striking was the miR-29 family, with miR-29a-3p and miR-29b-3p showing strong increases from newly hatched fry to adults and retaining negative associations with predicted targets after developmental stage was accounted for. miR-29 is a well-established regulator of extracellular-matrix genes, including collagens and other matrix components, and increasing miR-29 expression has been reported during postnatal tissue maturation^[19–21]^.

Together, these findings place the regulatory changes observed in *N. furzeri* within conserved miRNA programs that coordinate differentiation and tissue remodeling across vertebrates.

Importantly, the integrated analysis distinguished broad developmental co-variation from more persistent miRNA–target relationships. miR-200a, miR-141 and let-7b showed strong anti-correlated target trajectories across maturation but did not retain negative associations after developmental stage was accounted for, suggesting that these relationships primarily reflect coordinated developmental programs. In contrast, miR-29a, miR-29b, miR-34a, miR-15a and nfu-novel-017 retained repression-consistent target associations after stage adjustment.

Functional enrichment further linked anti-correlated target cohorts to developmental processes involving MAPK and FoxO signaling, cell adhesion, cytoskeletal organization, extracellular-matrix interactions and metabolism. These relationships remain predictive rather than mechanistic, but they identify specific miRNA–target modules for experimental follow-up.

The study also expands available miRNA annotation for *N. furzeri*, which remains absent from miRBase. Of 407 de novo candidates, 179 overlapped miRNAs previously predicted by Baumgart et al. (2017)^[23]^, providing independent sequence-level support for a substantial portion of the catalogue. Conservation analysis further showed that many apparently novel killifish miRNAs are evolutionarily conserved: 216 candidates shared seed sequences with mammalian miRNAs, whereas others showed more restricted conservation. A subset of developmentally dynamic miRNAs also overlapped families previously associated with ageing^[26]^, including miR-29 and let-7. This overlap is particularly interesting in a short-lived species with compressed maturation, but it should not be interpreted as an ageing signature because the present study follows a maturation series rather than an adult ageing series. Instead, it suggests that some regulatory trajectories associated with later life may begin during developmental maturation.

A major strength of this study is that miRNA and mRNA were measured from the same individual animals, enabling regulatory and target expression to be examined across matched biological samples. Sensitivity analyses further showed that the major target-enrichment patterns were maintained when adult males and females were analyzed separately and when alternative correlation approaches were used, supporting the robustness of the identified relationships.

Nevertheless, important limitations remain. Whole-organism profiling cannot fully distinguish cell-intrinsic regulatory changes from developmental shifts in tissue composition, although canonical tissue-marker miRNAs represented only a small fraction of the dynamic miRNA set. In addition, the three developmental stages capture major transitions across maturation but provide limited resolution for identifying transient regulatory events or precisely defining when individual miRNA–target relationships emerge. Finally, sequence-based target prediction and expression anti-correlation identify candidate regulatory relationships rather than direct miRNA– mRNA interactions. Future perturbation studies, reporter assays, and Argonaute-based approaches will therefore be important for validating specific interactions and establishing their functional roles during killifish development.

Overall, these findings identify post-hatch maturation in N. furzeri as a period of extensive coordinated miRNA and mRNA remodeling. Conserved regulators including let-7, miR-29, miR-200/141 and miR-34 connect the compressed killifish developmental trajectory to regulatory programs described across other animals, while stage-adjusted analyses identify a smaller set of candidate miRNAs with particularly robust repression-consistent target relationships. Together with the expanded species-specific miRNA catalogue, these data provide a framework for investigating how post-transcriptional regulation contributes to rapid vertebrate maturation. More broadly, the matched miRNA and mRNA datasets, developmental expression profiles, predicted miRNA–target relationships, and expanded miRNA annotations provide a useful resource for the killifish and developmental biology communities and a foundation for future mechanistic studies of post-transcriptional regulation during vertebrate development.

## Author Contributions

Z.A.: Investigation, Methodology, Formal analysis, Writing. C.H.: Methodology. D.K.: Conceptualization, Methodology, Supervision, Funding acquisition, Writing, Corresponding author.

## Ethics Statement

All animal experiments were conducted under Protocol No. DK10112024 approved by IACUC at North Carolina Central University.

## Data Availability

All data will be deposited on the GEO database and available to download.

## Acknowledgments

We acknowledge the Julius L. Chambers Biomedical/Biotechnology Research Institute (BBRI) at North Carolina Central University for providing research facilities and resources. We thank the BBRI faculty and staff and students for their support in this project, Vandana Veershetty, Evan E. Pittman, Dr. Derek Norford, Ms. Camilla Felton, Thomas J Broek, Dr. Qing Cheng, and Dr. Claudia Alberico. We are grateful to Dr. Anne Brunet and members of the Brunet Laboratory at Stanford University for their guidance in establishing the killifish facility and colony. We also thank Riley Galton and Robert Schnittker from the Sánchez Alvarado Laboratory at the Stowers Institute for all their support and expertise in guiding us for killifish embryo and juvenile husbandry, and Wouter Lanneau of Microbiotests for his assistance with killifish embryos. We also acknowledge the use of an AI agent for discussion and interpretation of the study findings.

Use of AI was done in August 2026 to assist with editing, refinement, organization and clarity of manuscript text, and development and troubleshooting of R code used for data visualization. It was not used to generate any of the main analysis or an underlying experimental data. All AI-assisted content, code, analyses, and interpretations were done according to established guidelines [27] and independently reviewed and verified by the authors, who take full responsibility for the accuracy and integrity of the manuscript.

## Figure legends

**Supplementary Figure S1. Stage-adjusted analysis distinguishes within-stage miRNA– target coupling from stage-driven associations.** (A) Stage-adjusted target-coupling scores for the 20 miRNAs identified by the initial target-enrichment analysis. miRNA and predicted-target expression were adjusted for developmental stage across the 10 matched animals before evaluating target associations. Negative z-scores indicate that predicted targets tended to show lower expression in individuals with higher expression of the corresponding miRNA. Red bars indicate miRNAs retaining significant negative target associations after stage adjustment (FDR<0.05). Five miRNAs—miR-29a, miR-34a, miR-29b, miR-15a and nfu-novel-017—met this criterion. (B) Comparison of across-stage target-enrichment and stage-adjusted target-coupling scores for the same 20 miRNAs. Red points indicate miRNAs retaining significant negative target associations after stage adjustment (FDR<0.05), whereas gray points indicate relationships primarily associated with the developmental-stage trajectory. miR-141, miR-200a and let-7b showed strong across-stage target enrichment but did not retain negative target coupling after stage adjustment, whereas miR-29a/b, miR-34a, miR-15a and nfu-novel-017 retained repression-consistent relationships after accounting for developmental stage.

**Supplementary Figure 2. Predicted target cohorts of stage-adjusted miRNAs shift toward reduced expression during maturation.** (A) Cumulative distributions of mRNA expression changes from newly hatched fish to adults for predicted target cohorts of miR-34a, miR-29a, miR-29b, miR-15a, and nfu-novel-017 compared with the genome-wide background (gray). Expression change is represented as the difference in variance-stabilized expression (VST) between adults and newly hatched fish, with negative values indicating decreased expression toward adulthood. A leftward shift relative to the genome-wide distribution indicates greater reduction in expression among predicted targets. (B) Median target-expression shift relative to the genome-wide background for miRNAs evaluated for repression-consistent target changes. Negative values indicate that predicted target cohorts decreased more strongly during maturation than the genomic background. Red indicates miRNAs that retained significant negative target associations after developmental stage was accounted for, whereas gray indicates miRNAs whose anti-correlated target relationships were primarily associated with the developmental-stage trajectory. Significance of target-cohort shifts relative to background was assessed using one-sided Mann–Whitney tests. ns, not significant; **P<0.01; ***P<0.001; ****P<0.0001.

