## Supplementary figures and images for "Coordinated microRNA–mRNA regulatory remodelling across the compressed maturation of the African killifish (*Nothobranchius furzeri*)"

### Supp Figures

#### Supp Figure 1

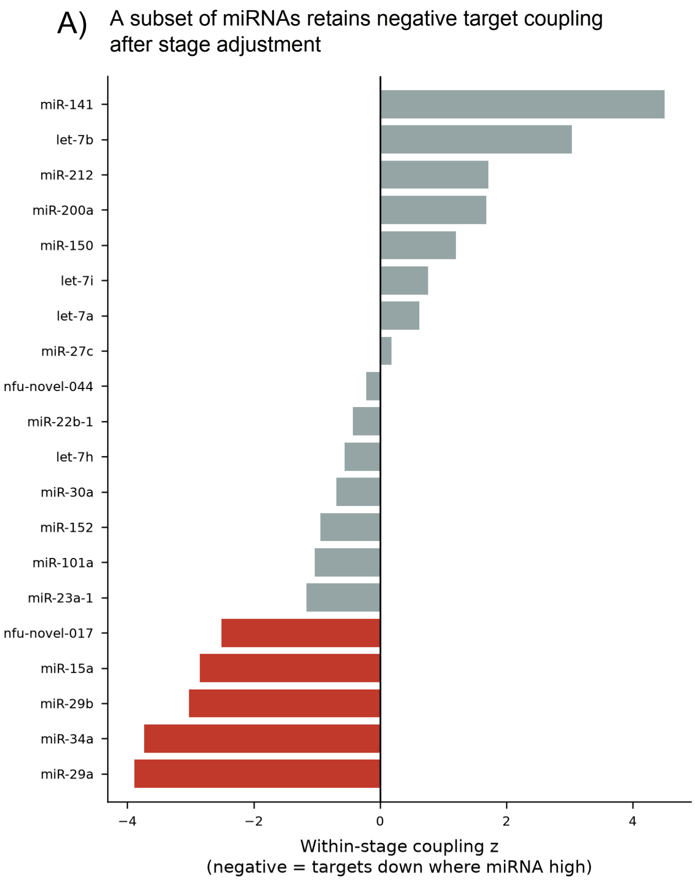


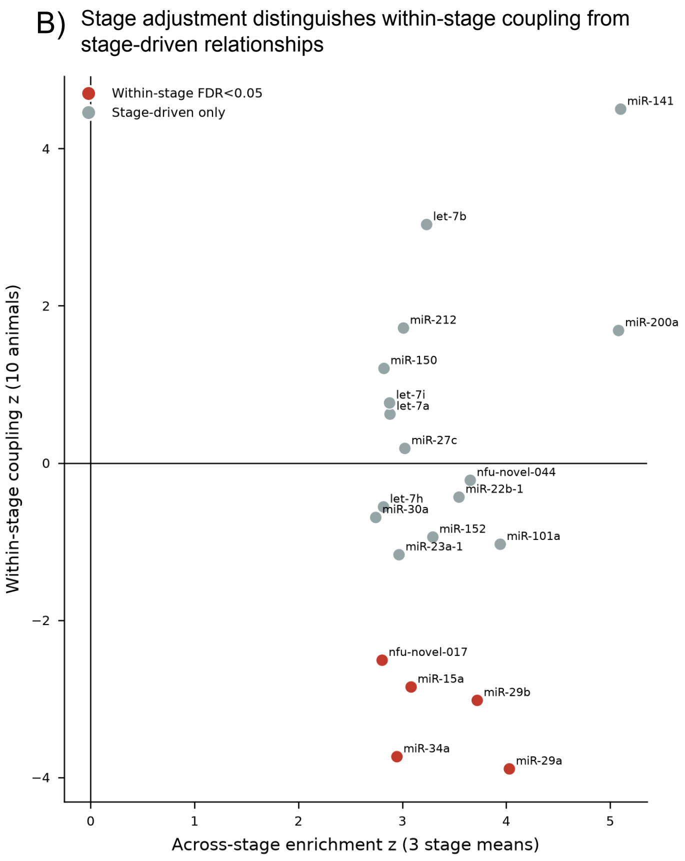

##### Supp Figure 2

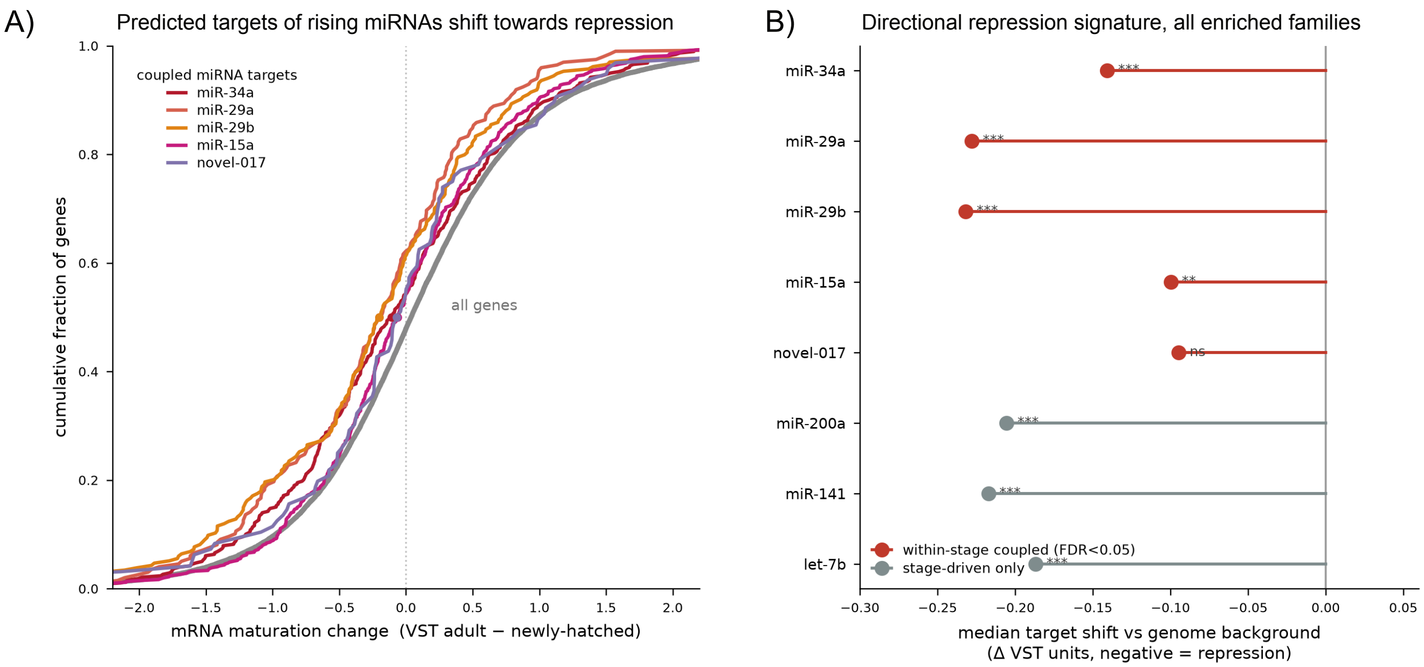
